# InSTaPath: Integrating Spatial Transcriptomics and histoPathology Images via Multimodal Topic Learning

**DOI:** 10.64898/2026.03.16.712067

**Authors:** Weiyi Xiao, Hegang Chen, Adrien Osakwe, Qihuang Zhang, Yue Li

## Abstract

Spatial transcriptomic (ST) technologies enable the measurement of gene expression directly within tissue sections while preserving spatial context. Many ST platforms additionally generate paired histological images alongside spatially resolved transcriptomic profiles. However, most existing computational approaches only incorporate histology images as auxiliary features in representation learning models and typically produce latent embeddings that are difficult to interpret. We present InSTaPath (<u>Int</u>egrating <u>S</u>patial <u>T</u>ranscriptomics <u>a</u>nd histo<u>Path</u>ology images), a multimodal topic modeling framework that links transcriptional programs with tissue morphology. InSTaPath converts token-level embeddings extracted from pretrained histology foundation models into discrete image words through vector quantization, enabling histological morphology to be represented in a count-based form analogous to gene expression. InSTaPath then jointly analyzes image-word and gene expression counts to infer shared latent topics that are interpretable through both topic–gene and topic–image-word associations. Across multiple ST datasets, InSTaPath improves spatial domain identification and uncovers biologically meaningful relationships between gene programs and tissue morphology through pathway enrichment and *in silico* perturbation analyses.

## Introduction

Spatial transcriptomics (ST) technologies enable measurement of gene expression while preserving the spatial organization of tissues (Ståhl et al., 2016). Many platforms (e.g., 10x Visium) generate paired multimodal data consisting of a histological Hematoxylin and Eosin (H&E) stained image of the tissue section together with spatially resolved gene expression profiles at discrete capture spots. These data provide an opportunity to connect transcriptional programs with tissue morphology. However, in many existing analysis pipelines, histology images are primarily used for visualization or manual annotation, whereas downstream computational analyses rely mainly on gene expression. Consequently, they fail to capture interactions between tissue architecture and transcriptional programs.

Several recent studies have sought to integrate histology with gene expression using deep learning approaches, which can be broadly divided into two categories. The first comprises graph- and convolution-based methods that extract image features from patches surrounding each capture spot and combine them with gene expression to infer spatial representations (e.g., SpaGCN (Hu et al., 2021), DeepST (Xu et al., 2022)). The second includes foundation model-based approaches that leverage large-scale representation learning to align image and molecular modalities. These include contrastive learning methods inspired by CLIP (Radford et al., 2021) (e.g., OmiCLIP (Chen et al., 2025)) or masked modeling approaches such as STPath (Huang et al., 2025). Although these methods demonstrate the benefits of integrating histology and gene expression data, they do not explicitly model shared latent programs linking transcriptional and morphological signals, thereby limiting model interpretability.

Topic modeling provides a natural framework for interpretable ST modeling, as it can represent each ST spot as a mixture of latent programs, where each program defines an interpretable distribution over features (e.g., genes) (Osakwe et al., 2025; Zhong et al., 2024). However, extending topic models to histology is challenging, as the data are represented by high-dimensional continuous pixel values rather than discrete count data.

Here we introduce InSTaPath (Integrating Spatial Transcriptomics and Histopathology images), a framework that jointly models gene expression and histological morphology using multimodal topic modeling (Fig.1). InSTaPath first converts histology images into a discrete representation by extracting token-level embeddings from a pretrained histology foundation model and mapping them to *image words* through vector quantization. These image words are then aggregated within each ST capture spot and jointly modeled alongside gene expression counts in a shared topic model framework. This design enables each latent topic to be interpreted simultaneously through gene- and image-based associations, thereby linking transcriptional programs with morphological tissue patterns in a unified probabilistic framework. In addition, as the image words are derived from token-level features extracted from whole-slide histology images, the framework can incorporate broader morphological context beyond spot-centered patches measured by the ST platform. The resulting topic representations support multiple downstream analyses, including characterization of spatial tissue programs, biological interpretation through gene set enrichment, and *in silico* perturbation analyses that predict how changes in gene expression alter morphology-associated signals.

**Figure 1.**
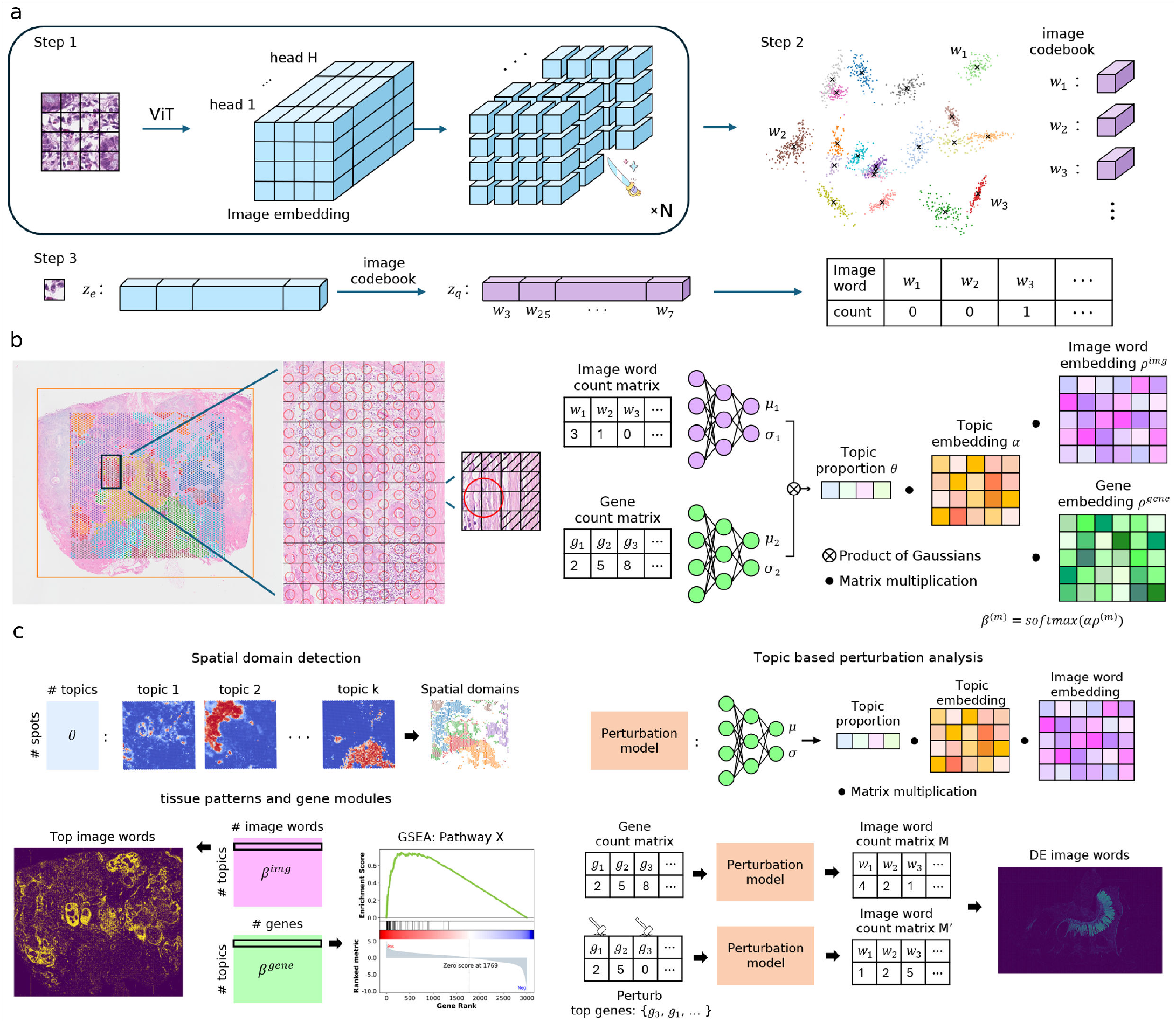
Overview of InSTaPath and downstream analyses. **(a)** Construction of image words from histology. Step 1: extraction of token-level image embeddings. Step 2: codebook generation. Step 3: vector quantization to obtain token-level image word count vectors. **(b)** Integration with spatial transcriptomics. Token-level image word counts on the whole-slide image (WSI) are aggregated into spot-level counts based on the spatial footprint of each spot. In the illustration, each patch contains 4 × 4 tokens and the selected spot covers 8 tokens, whose counts are summed to obtain the spot-level image representation. Paired gene counts and image word counts are then jointly modeled to infer shared latent topics linking molecular programs and tissue morphology. **(c)** Downstream analyses based on the learned topic representations. The spot–topic distribution (*θ*) is used for spatial domain detection. The topic–image word distribution (*β*^*img*^) identifies top image words, which can be visualized on the WSI to reveal representative tissue patterns. The topic–gene distribution (*β*^*gene*^) ranks genes within each topic and enables pathway enrichment analysis such as GSEA. Finally, *in silico* perturbation analysis uses the trained gene encoder and image decoder to reconstruct image features from gene expression and assess the causal influence of genes on image words.

## Materials and methods

### InSTaPath workflow

InSTaPath enables an integrative analysis of paired histology images and ST data in three stages. In Stage 1, a pretrained foundation model is used to extract token-level embeddings from whole-slide histology images. These embeddings are then discretized via vector quantization to produce count-based image-word representations (Fig. 1a). In Stage 2, paired image-word and gene expression counts are jointly modeled across spots using multimodal topic modeling to infer latent topics linking morphology and gene expression (Fig. 1b). Finally, in Stage 3, we apply InSTaPath for spatial domain detection, gene program inference, and *in silico* perturbation analysis (Fig. 1c). The following sections provide details for each stage.

#### Stage 1 - Image embedding discretization

To construct a discrete representation of histology images suitable for topic modeling, InSTaPath converts continuous image embeddings into count-based image words. This procedure consists of three steps: (1) extraction of token-level image embeddings from whole-slide histology images (WSIs), (2) construction of a visual codebook, and (3) vector quantization of token embeddings to obtain image-word counts (Fig. 1a).

##### Step 1: Token-level image embedding extraction

WSIs are divided into smaller image tiles, each of which is processed by a Vision Transformer (ViT) to extract token-level embeddings. We use the pretrained histology foundation model UNI (Chen et al., 2024) as the backbone encoder. UNI employs a ViT-Giant architecture (`vit giant patch14 224`) configured to process 224×224 pixel patches at a resolution of 0.5 *µ*m*/*px with a token size of 14 × 14 pixels. The resulting embeddings (dimension 1536) are used for discretization.

##### Step 2: Codebook construction

To discretize the continuous embeddings, each token embedding is partitioned into sub-vectors with dimensionality matching the attention head dimension (e.g., 64). Sub-vector lengths equal to either half or the full head dimension generally provide strong performance in downstream tasks. We then apply mini-batch *k*-means clustering to the collection of sub-vectors extracted from all image tiles. The resulting cluster centers define the entries of a visual codebook for vector quantization.

##### Step 3: Vector quantization and image-word representation

Inspired by the vector quantization variational autoencoder (VQ-VAE, Van Den Oord et al. (2017)), each cluster center is treated as an image word, and each sub-vector is assigned to its nearest codebook entry. Through this process, each token embedding is converted into a count vector of codebook entries. In practice, we use a codebook of 512 image words with 64-dimensional sub-vectors, resulting in a 512-dimensional count vector for each token embedding. This discrete image-word representation enables the application of topic modeling to histological images and allows image features to be jointly modeled with gene expression in a multimodal framework.

Additional analyses show that the ViT-VQ representation retains comparable discriminative information to the original UNI embeddings and is robust to the ordering of feature dimensions (Supplementary Section S5).

#### Stage 2 - Topic modeling of paired image and gene expression

We treat each spatial location (i.e., an ST spot) as a multimodal “document” with two modalities: histological image features and gene expression. InSTaPath identifies co-occurring image words and genes across spots to discover biologically meaningful topics (Fig. 1b). To jointly model these modalities, InSTaPath employs modality-specific VAE encoders, each implemented as a two-layer fully connected neural network that maps input to a latent topic representation. A product-of-Gaussians mechanism (Zhou et al., 2023) then integrates the modality-specific latent variables into a unified spot-level topic distribution, denoted by *θ* ∈ ℝ^*N*×*K*^, where *N* and *K* are the number of spatial spots and latent topics, respectively, and each row of *θ* sums to one. Given *θ*, the decoder reconstructs both modalities through modality-specific feature embeddings. Specifically, InSTaPath uses a shared topic embedding matrix *α* ∈ ℝ^*K*×*D*^ in a common *D*-dimensional latent space. For each modality *m* ∈ {gene, img}, a modality-specific feature embedding matrix 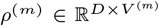 maps genes or image words into this shared space, where *V* ^(*m*)^ is the vocabulary size of modality *m*. The topic–feature distributions are defined as *β*^(*m*)^ = softmax(*αρ*^(*m*)^), where 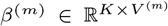 and each row represents a probability distribution over features for one topic. In practice, we use the top 3000 highly variable genes and the top 256 highly variable image words. Details are provided in Section S2.

#### Stage 3 - Integrative image-ST analyses

InSTaPath leverages several interpretable outputs from Stage 2 for downstream analysis of ST data, including (1) a spot-level topic representation (*θ*) summarizing the mixture of tissue programs at each spatial location, (2) topic–gene association scores (*β*^*gene*^) ranking genes by their relevance to each topic, and (3) topic–image-word associations (*β*^*img*^) enabling visualization of morphological patterns linked to each topic. These quantities provide a unified representation of multimodal tissue structure and support multiple downstream analyses. In this study, we use the spot–topic representations for spatial domain detection, the topic–gene scores for gene set enrichment analysis, and the gene-to-image reconstruction framework for *in silico* perturbation experiments that evaluate how changes in gene expression influence predicted histological patterns.

##### Spatial domain detection

The inferred spot–topic proportions, *θ*, provide a low-dimensional multimodal representation of each spatial location. These topic mixtures capture co-occurring transcriptional and morphological patterns and serve as features for spatial domain detection. We performed clustering on *θ* using the mclust R package (Scrucca et al., 2023), where the number of clusters was specified based on the known number of tissue regions in the dataset. Clustering performance was evaluated using Adjusted Rand Index (ARI), Normalized Mutual Information (NMI), and Average Silhouette Width (ASW) (Section S3).

##### Visualization of topic-associated tissue patterns

To visualize morphological patterns associated with individual topics, we plotted the spatial distribution of the top five image words for each topic based on their topic probabilities in *β*^(img)^. We computed the occurrence of these image words across all tokens in the WSIs and summed their counts at each token location to obtain a topic-specific image-word expression value. The resulting heatmaps highlight regions where the top image words are enriched, enabling visualization of tissue structures associated with each topic. These maps can be compared with the underlying histology image or with feature maps derived from the vision backbone to interpret the morphological patterns captured by the learned topics.

##### Gene set enrichment analysis for the inferred topics

To interpret the biological functions associated with each latent topic, we performed gene set enrichment analysis (GSEA) using the topic–gene scores *β*^(gene)^ learned by InSTaPath. GSEA was performed using the gseapy Python package (Fang et al., 2023). For each topic, genes were sorted in descending order of their raw scores and supplied to the `pre-rank` procedure. Enrichment was evaluated against several human MSigDB collections, including H (Hallmark), C2 (Curated), C4 (Computational), C5 (Gene Ontology), and C6 (Oncogenic signatures).

##### *In silico* perturbation

To investigate how transcriptional programs influence histological patterns, we performed *in silico* gene perturbation experiments using the trained InSTaPath model. Gene expression profiles are propagated through the gene encoder and image decoder to reconstruct predicted image-word distributions. Given a gene expression matrix *X*^(gene)^, the gene encoder first produces the spot–topic representation *θ*, which is then used by the image decoder to impute image-word probabilities as 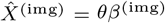. To simulate gene perturbations, we selected the top-ranked genes associated with each topic based on the topic–gene scores and set their expression values to zero across all spatial locations. The perturbed gene expression matrix was then passed through the trained encoder–decoder pipeline to produce image-word predictions, reflecting how gene perturbation affects histological signals (i.e., endophenotypes).

To quantify the effect of transcriptional perturbations on histological signals, we trained a logistic regression classifier on the reconstructed image-word features to distinguish a tissue of interest from other regions based on spot-level annotations (e.g., tumor vs. non-tumor). The reconstructed image-word profiles obtained after perturbation were evaluated using the trained classifier to obtain updated probabilities. The perturbation effect was summarized as the mean change in predicted probability for the target tissue region relative to the original predictions. By progressively perturbing increasing numbers of top-ranked genes associated with each topic, we quantified how predicted histological signals respond to transcriptional perturbations linked to different tissue programs. The same procedure can be applied to diverse tissue programs identified by InSTaPath (e.g., tumor, smooth muscle, fibroblast, or immune-associated regions). Qualitatively, we visualized the spatial effects of perturbations by comparing reconstructed image-word signals before and after perturbation. We also identified image words that exhibited the largest changes between perturbed and unperturbed conditions and mapped their spatial distributions back onto the WSI, enabling visualization of perturbation-induced morphological changes beyond the ST captured area.

## Results

### InSTaPath learns visual topics from pathology images

To demonstrate that the image-word representation derived from discretized histology embeddings captures meaningful tissue patterns, we analyzed the CRC-100k colorectal cancer dataset (Kather et al., 2019), consisting of 100,000 non-overlapping image patches of size 224 × 224 pixels extracted from H&E-stained histological slides of colorectal cancer and normal tissue from 86 patients and annotated with nine tissue classes. Using the image-only mode of InSTaPath, each image patch is treated as a document composed of image words derived from discretized embeddings. We compared embeddings derived from the pretrained histology foundation model (UNI), the discretized image-word representation (ViT-VQ), and the patch–topic embedding learned by InSTaPath using *K* = 50 topics (Fig. 2a). Raw patch embeddings extracted from UNI exhibit partial separation across tissue classes but often form scattered clusters, possibly reflecting patient-specific variation or batch effects. After vector quantization using a codebook of 512 image words, the representation became more discrete, resulting in clearer separation between clusters and stronger alignment with histological classes. Applying InSTaPath further enhanced the semantic organization of tissue patterns, producing tightly grouped clusters of patches that shared strong topic associations and corresponded more clearly to ground truth classes.

**Figure 2.**
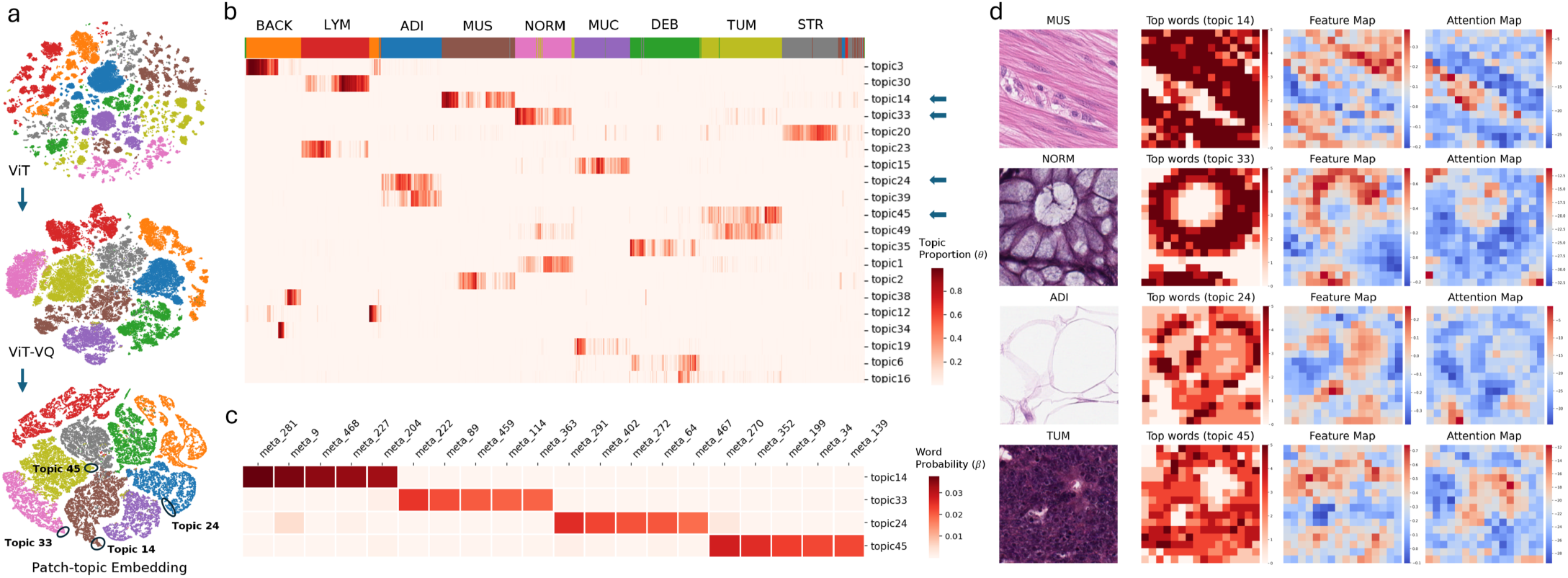
Analysis of CRC-100k colorectal cancer pathology imaging data using InSTaPath with only the image-component enabled. **(a)** t-SNE visualization of patch features from an image foundation model (e.g., UNI), vector-quantized patch features, and patch–topic embeddings. The circled regions highlight the top 100 patches for four representative topics. **(b)** Clustermap of the patch–topic distribution *θ*, showing the top 20 topics. In the clustermap, each column represents an image patch and each row corresponds to a topic, with color intensity indicating the topic proportion. **(c)** Heatmap of the topic–word distribution for the four selected topics from part (b). **(d)** For each selected image, four subplots are shown: the image patch, the spatial distribution of top-word counts, the feature map, and the attention map. The patch topic is determined as the topic with the largest topic proportion. The total occurrence count of the top-5 image words from that topic is summed for each token and then plotted spatially. The feature map shows the mean value of the token embeddings. The attention map shows the attention from the [*CLS*] token. Abbreviations: ADI: adipose; BACK: background; DEB: debris; LYM: lymphocytes; MUC: mucus; MUS: smooth muscle; NORM: normal colon mucosa; STR: cancer-associated stroma; TUM: colorectal adenocarcinoma epithelium.

Visualizing the learned patch–topic distribution *θ* (Fig. 2b) revealed morphology-specific topics, with high topic proportions concentrated in subsets of patches that predominantly map to the same histological class. Quantitatively, InSTaPath achieves a topic diversity of 0.77 and a topic coherence of 0.13 (Section S3). The high topic diversity indicates that InSTaPath identifies mostly distinct top image words for each topic (Fig. 2c). To demonstrate the biological relevance of these top image words, we visualized the spatial distribution of top-word occurrences (Fig. 2d). Compared with the feature and attention maps derived from the ViT model, the top-word occurrence maps align more closely with the underlying tissue structures. For example, topics assigned to muscle and normal glandular structures emphasize elongated fiber patterns and glandular lumens, respectively, whereas tumor-associated topics focus on densely packed cellular regions. These results are robust to the choice of topic number, with similar visual patterns and topic quality observed for *K* values between 10 and 50.

### InSTaPath detects spatial domains from breast cancer data

Motivated by the performance of InSTaPath on imaging data, we applied InSTaPath with *K* = 10 topics to 10x Visium breast cancer data (Janesick et al., 2023), which contains both gene expression and histology data. To illustrate that integrating both modalities may yield higher performance in learning spot representations, we compared InSTaPath with single-modality baselines. InSTaPath produced spatial clusters that more closely matched the ground-truth tissue annotations, achieving higher clustering accuracy (ARI = 0.82) than using gene expression (ARI = 0.60) or histology images (ARI = 0.47) alone (Fig. 3a). InSTaPath also outperformed the GNN-based topic modeling method STAMP and the foundation model-based approach OmiCLIP in terms of ARI, NMI, and ASW for the spatial domain detection task (Table 1).

**Table 1.** Clustering performance comparison.

| Method | ARI | ASW | NMI |
| --- | --- | --- | --- |
| InSTaPath (gene+img) | **0.82** | **0.58** | **0.86** |
| OmiCLIP (gene) | 0.80 | 0.20 | 0.80 |
| STAMP (gene) | 0.77 | 0.42 | 0.83 |
| OmiCLIP (gene+img) | 0.70 | 0.19 | 0.76 |
| InSTaPath (gene) | 0.60 | 0.42 | 0.65 |
| InSTaPath (img) | 0.47 | 0.21 | 0.54 |
| OmiCLIP (img) | 0.43 | 0.09 | 0.46 |

ARI: Adjusted Rand Index; ASW: Average Silhouette Width; NMI: Normalized Mutual Information. Note that we only considered spots with single labels for evaluation, namely DCIS#1, DCIS#2, invasive, stromal, adipocytes, and immune. All metrics are reported as higher-is-better. Details of the benchmarked methods are described in Supplementary Section S4.

**Figure 3.**
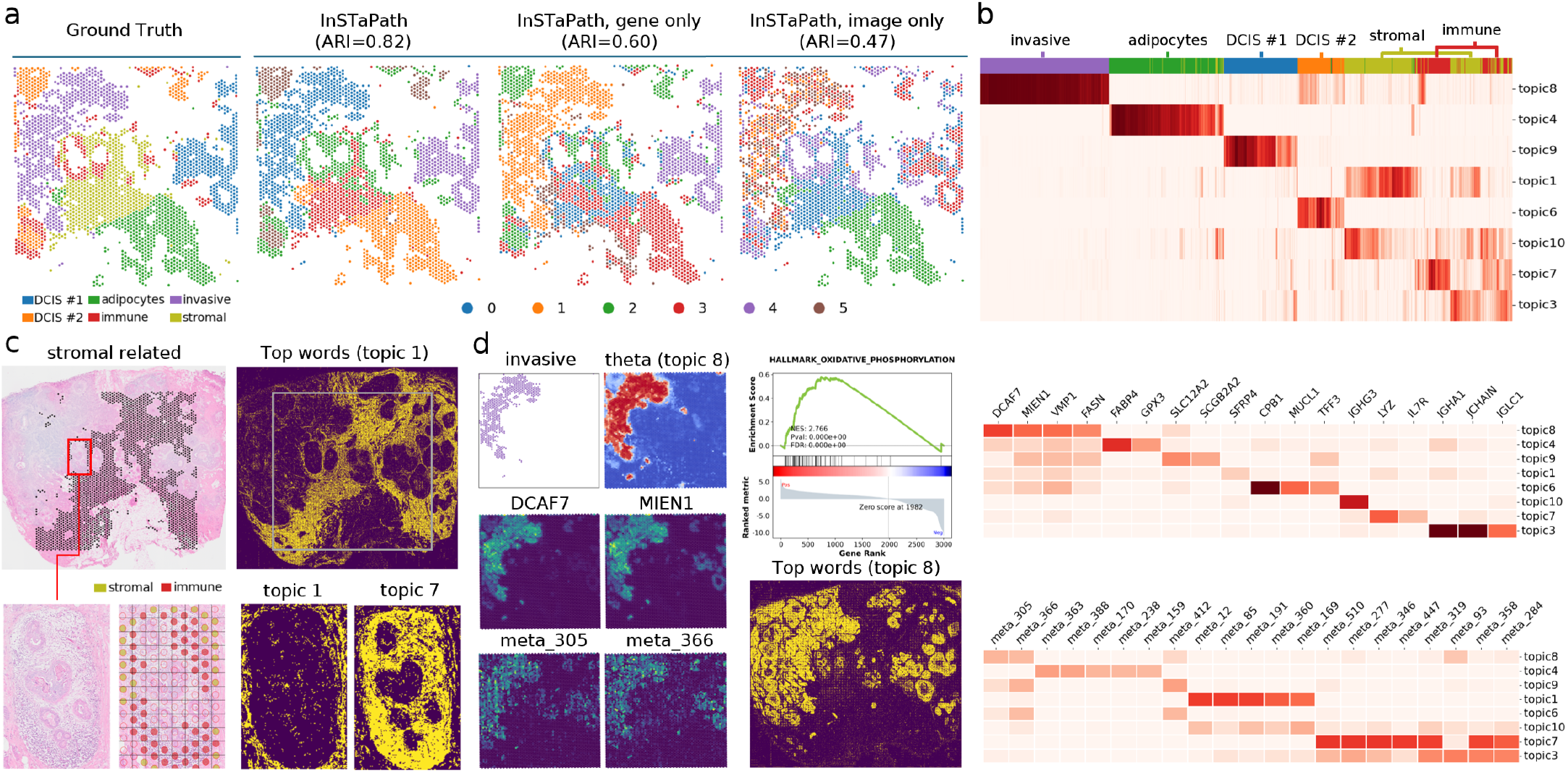
Analyzing 10x Visium breast cancer data with InSTaPath by integrating gene expression and histology. **(a)** Spatial clustering comparison across models. Using six ground-truth spot annotations (DCIS#1, DCIS#2, invasive, stromal, adipocytes, and immune), *InSTaPath* achieves improved clustering accuracy compared with single-modality baselines using gene expression or histology features. **(b)** Top: Spatial distributions of representative InSTaPath topics (*θ*); Bottom: Heatmaps of top topic-specific features (*β*^(img)^, *β*^(gene)^). **(c)** Histology-informed separation of stromal and immune regions. Topic 1 corresponds to stromal regions and topic 7 to immune regions. A zoomed region highlights that topic-specific image words capture distinct morphological features. **(d)** Alignment of gene expression and histology signals and their applications. The left panels show the aligned spatial distributions of representative genes and image words; the right panels show GSEA results and whole-slide distributions of the corresponding top image words.

The inferred spot–topic distribution showed clear correspondence with tissue structures, with individual topics capturing distinct anatomical regions. For example, topic 8 localized to invasive tumor regions and topic 4 corresponded to adipocyte-rich areas (Fig. 3b). The gene and image modalities provide complementary information for spatial domain identification. Gene-only methods can separate transcriptionally distinct regions such as invasive tumor and DCIS subtypes, but often fail to distinguish morphologically defined compartments such as stromal and immune regions. In contrast, histology-based features better capture these morphological differences but are less effective at separating transcriptionally similar tumor regions. Using a zoomed region containing mixed stromal and immune spots (red box in Fig. 3c), we show that the top image words associated with topic 1 and topic 7 highlight distinct morphological patterns corresponding to stromal and immune compartments. Gene-only methods such as STAMP (Zhong et al., 2024) failed to separate these regions effectively, whereas methods incorporating histology information, including InSTaPath and OmiCLIP (Chen et al., 2025), achieved clearer separation (Fig. S3). By jointly modeling both gene expression and histological image features, InSTaPath integrates these complementary signals and produces spatial clusters that better match the underlying tissue organization. In addition, the learned image words enable visualization of topic-associated tissue patterns across the entire histology slide beyond the spatial transcriptomics capture area (Fig. 3c, right column).

### InSTaPath links transcriptome to WSI-derived phenotypes

Moreover, InSTaPath captures aligned gene and histological signals within individual topics (Fig. 3d). The top genes and top image words for each topic show similar spatial distributions. To understand the biological meaning of the tissue patterns associated with each topic, we performed gene set enrichment analysis using the gene topic scores. For example, the invasive-associated topic 8 showed enrichment for oxidative phosphorylation pathways (Fig. 3d), the adipocyte-associated topic 4 was enriched for adipogenesis signatures, and the immune-associated topic 7 showed enrichment for T-lymphocyte differentiation (Fig. S4). Together, these results demonstrate that InSTaPath effectively integrates spatial transcriptomic and histological information to uncover biologically meaningful spatial programs.

### InSTaPath enables *in silico* gene perturbation on pathology

To further demonstrate InSTaPath, we tested it on Visium HD colorectal cancer data (Oliveira et al., 2024) using *K* = 10 topics. It identified topics that correspond well to the annotation labels. For example, topic 1 corresponds to the smooth muscle region, topic 3 to goblet cells and enterocytes, and topic 6 to tumor regions (Fig. S5b). An advantage of InSTaPath is its ability to identify genes and image words that co-occur. Using the tumor-associated topic 6 as an example, we zoomed in on a region of interest (ROI) (Fig. S5c). Within the ROI, the spatial distributions of representative top image words and genes are highly concordant with tumor morphology. When visualized across the full histology image, the top image words for topic 6 consistently highlight the tumor-associated tissue architecture (Fig. S5d).

Next, we quantified how transcriptional perturbations influence the histologic signals captured by the model. Following the *in silico* perturbation procedure described in the Methods, we evaluated the reconstructed image-word representations using a logistic regression classifier trained to distinguish tumor versus non-tumor regions.

Perturbing the top five genes for each topic revealed that tumor-associated topics produced the largest decreases in predicted tumor probability, particularly for topics 4, 5, and 6 (Fig. 4b).

**Figure 4.**
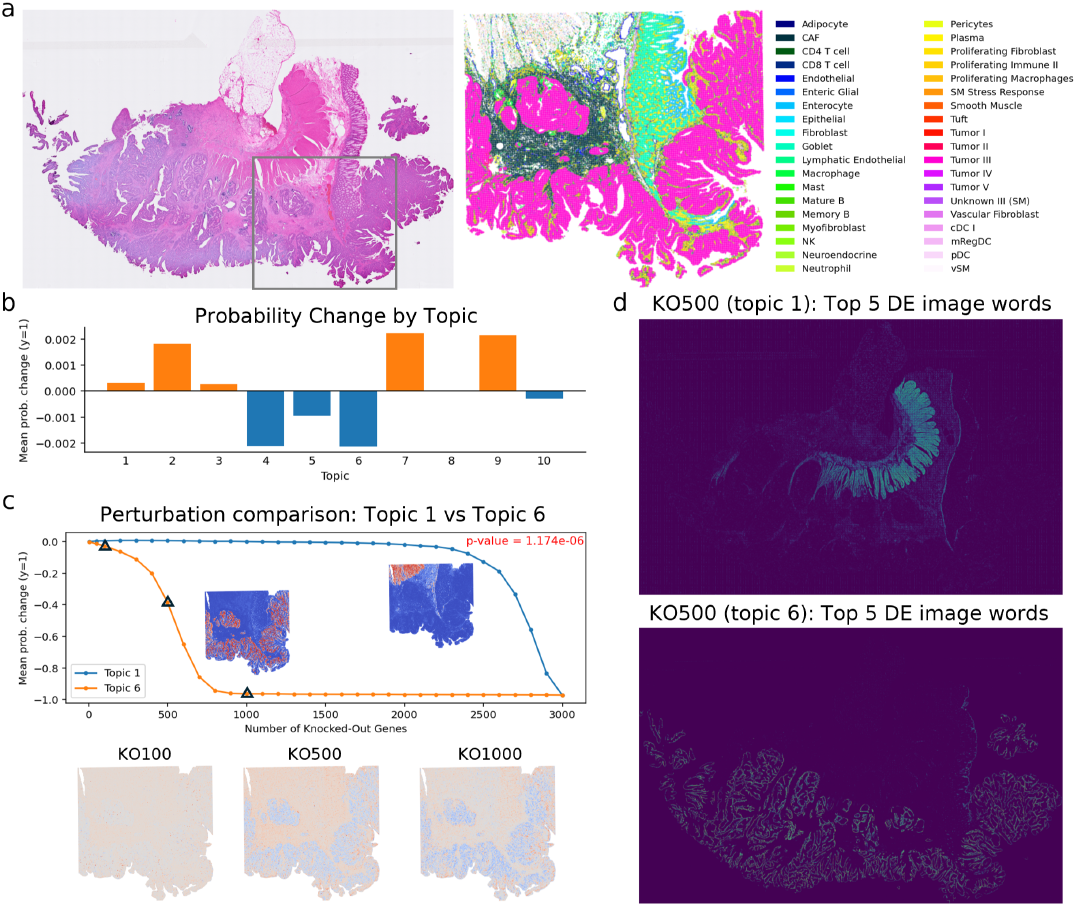
In silico gene perturbation analysis. (**a**) Whole-slide histology image (left) and corresponding cell-type annotation map (right). The gray box indicates the region used for spatial transcriptomics profiling. (**b**) *In silico* gene knockout analysis. A logistic regression classifier trained on image words predicts tumor versus non-tumor spots. After knocking out the top five genes associated with each topic, we measure the mean change in tumor prediction probability for each topic. (**c**) Comparison of perturbation effects between topic 6 (tumor) and topic 1 (smooth muscle). P-values were calculated using the Wilcoxon rank-sum test. The bottom panel shows the change in the top image words for topic 6 after knocking out its top 100, 500, and 1000 genes. (**d**) Spatial distribution of the top five differentially expressed (DE) image words after gene perturbation. Image words with the largest log-fold changes between perturbed (KO500) and original reconstructions were identified for each topic, and their occurrence counts were visualized across the WSI.

We further examined topic-specific perturbation effects by progressively removing increasing numbers of top-ranked genes (e.g., the top 5, 50, and 100 genes) from different topics. For tumor-associated topic 6, the predicted tumor probability decreased rapidly as more genes were removed, whereas the smooth muscle-associated topic 1 showed a substantially slower decline. Consistently, removing more top genes from topic 6 reduced the occurrence of the corresponding tumor-associated image words within tumor regions (Fig. 4c). To visualize the spatial impact of these perturbations, we identified the image words showing the largest differences between perturbed and unperturbed conditions and mapped their spatial distributions across the whole-slide image. For topic 1, knocking out the top 500 genes produced differentially expressed image words that visually align with the smooth muscle region on the whole-slide image. For topic 6, the top differentially expressed image words correspond closely to tumor regions across the tissue section and capture fine-grained structural features (Fig. 4d).

## Conclusion

In this work, we presented InSTaPath, a multimodal framework that integrates spatial transcriptomics and histopathology through interpretable topic modeling. By converting continuous histology embeddings from pretrained foundation models into discrete image words and jointly modeling them with gene expression counts, InSTaPath identifies latent topics that capture shared morphological and transcriptional structure across spatial locations. Each topic is interpretable through both topic–gene and topic–image-word associations.

In breast and colorectal cancer case studies, InSTaPath uncovers spatially coherent tissue programs and links gene expression with tumor microenvironment features. Image words derived from token-level features of whole-slide histology images enable visualization of topic-associated morphology beyond spot-centered patches and outside the spatial transcriptomics capture region. Perturbation analyses of gene–morphology relationships further highlight candidate marker genes in cancer-associated topics. Future work will extend the framework to additional imaging modalities beyond H&E-stained histology and improve robustness to variability in staining, resolution, and tissue preparation.

## Acknowledgment

This work is supported by New Frontier Research Fund Exploration (NFRFE-2024-00691). We thank Li and Zhang lab members for helpful discussion.

## Data availability

We analyzed the following datasets: (1) histological images of human colorectal cancer and healthy tissue, CRC-100K (https://zenodo.org/records/1214456); (2) 10x Genomics human breast cancer Visium and H&E image data (https://www.10xgenomics.com/products/xenium-in-situ/preview-dataset-human-breast); (3) 10x Genomics human colorectal cancer and colon non-diseased Visium HD and H&E image data, Sample P2 CRC (https://www.10xgenomics.com/products/visium-hd-spatial-gene-expression/dataset-human-crc). The source code of InSTaPath is available in GitHub, at (https://github.com/xwymary/InSTaPath).

