## Supplementary Materials for "InSTaPath: Integrating Spatial Transcriptomics and histoPathology Images via Multimodal Topic Learning"

#### S1 Dataset description

**CRC-100k and CRC-7k**: This dataset is curated as training and testing sets for training vision foundation models to learn visual concepts in histopathology. CRC-100k ("NCT-CRC-HE-100K-NONORM") consists of 100,000 non-overlapping image patches across nine classes, extracted from hematoxylin and eosin (H&E)-stained histological images of human colorectal cancer (CRC) and normal tissue from formalin-fixed paraffin-embedded (FFPE) samples. These image patches were manually extracted from $N = 86$ patients. CRC-7k consists of 7,180 image patches across the same nine classes from $N = 50$ patients, with no patient overlap with NCT-CRC-HE-100K-NONORM. The nine tissue classes are: Adipose (ADI), Background (BACK), Debris (DEB), Lymphocytes (LYM), Mucus (MUC), Smooth muscle (MUS), Normal colon mucosa (NORM), Cancer-associated stroma (STR), and Colorectal adenocarcinoma epithelium (TUM). Following the ViT tokenization scheme, each image patch contains $16 \times 16 = 256$ image tokens.

**10x Genomics human breast cancer Visium and H&E image data:** This dataset contains 4,992 spatial spots and 18,074 genes. Each spot covers approximately 55 image tokens on average. The dataset includes 11 annotated categories (spot counts in parentheses): adipocytes (505), DCIS #1 (344), DCIS #2 (210), immune (263), invasive (599), mixed (438), mixed/invasive (369), myoepithelial/stromal/immune (110), stromal (555), stromal/endothelial (1,005), and stromal/endothelial/immune (594). All spots are used for model training, but only the six categories corresponding to single spot types are used for evaluation, namely DCIS #1, DCIS #2, adipocytes, immune, invasive, and stromal.

**10x Genomics human colorectal cancer and colon non-diseased Visium HD and H&E image data, Sample P2 CRC:** This dataset contains 545,613 spatial spots and 18,074 genes. Each spot covers approximately 1–3 image tokens. The dataset includes 38 annotated cell-type and tissue categories (spot counts in parentheses): Adipocyte (276), CAF (59,317), CD4 T cell (3,711), CD8 T cell (2,447), cDC I (726), Endothelial (18,040), Enteric Glial (857), Enterocyte (10,381), Epithelial (171), Fibroblast (3,363), Goblet (30,300), Lymphatic Endothelial (1,935), Macrophage (15,075), Mast (417), Mature B (213), Memory B (29), mRegDC (476), Myofibroblast (6,107), Neutrophil (4,641), Neuroendocrine (343), NK (12), pDC (105), Pericytes (6,011), Plasma (12,066), Proliferating Fibroblast (9,158), Proliferating Immune II (3,660), Proliferating Macrophages (9,243), SM Stress Response (60), Smooth Muscle (1,345), Tuft (88), Tumor I (258), Tumor II (1,537), Tumor III (213,672), Tumor IV (141), Tumor V (249), Unknown III (SM) (1,112), Vascular Fibroblast (62), and vSM (13,225).

#### S2 Multimodal embedded topic model

We model each spatial transcriptomics (ST) spot as a multimodal "document" containing two modalities: image-word counts derived from histology and gene-expression counts measured by the ST assay. InSTaPath maps each modality to latent topic representations using modality-specific VAE encoders and integrates them through a product-of-Gaussians mechanism to obtain a unified spot-level topic distribution. Given the inferred spot–topic distribution $\theta$, the decoder reconstructs both modalities through modality-specific feature embeddings. The matrix $\theta \in \mathbb{R}^{N \times K}$ represents the topic proportions for all spatial spots, where $N$ is the number of spots and $K$ is the number of topics; each row of $\theta$ sums to one because it represents a probability distribution over topics. InSTaPath uses a shared topic embedding matrix $\alpha \in \mathbb{R}^{K \times D}$ that encodes the latent topics in a common $D$-dimensional embedding space. For each modality $m \in \{\text{gene}, \text{img}\}$, a modality-specific feature embedding matrix $\rho^{(m)} \in \mathbb{R}^{D \times V^{(m)}}$ maps the features (genes or image words) into this shared space, where $V^{(m)}$ denotes the vocabulary size of modality $m$. The topic–feature distributions are then obtained as

$$\beta^{(m)} = \text{softmax}(\alpha \rho^{(m)}),$$

where $\beta^{(m)} \in \mathbb{R}^{K \times V^{(m)}}$ represents the topic–feature associations and each row sums to one. In practice, the gene vocabulary consists of the top 3000 highly variable genes, while the image vocabulary consists of the top 256 highly variable image words. This factorized parameterization allows the model to share the topic structure across modalities while maintaining modality-specific feature representations.

The generative process of InSTaPath is as follows:

1. For spot $s$, draw the topic proportions $\theta_s$ over the $K$ topics: $\theta_s \sim \mathcal{LN}(0, I_K)$, where $\mathcal{LN}(\cdot)$ denotes the logistic-normal distribution. Sampling from this distribution proceeds as

$$\delta_s \sim \mathcal{N}(0, I_K), \quad \theta_s = \text{softmax}(\delta_s), \quad \delta_s, \theta_s \in \mathbb{R}^K.$$

2. For each modality $m \in \{\text{img}, \text{gene}\}$ and each word $n$ in spot $s$:
   a. Draw a topic assignment $z_{sn} \sim \text{Cat}(\theta_s)$.
   b. Draw the word $w_{sn}^{(m)} \sim \text{softmax}\left(\beta_{z_{sn}}^{(m)}\right)$, where $\beta_{z_{sn}}^{(m)} = \rho^{(m)\top} \alpha_{z_{sn}}$. Here, $\rho^{(m)}$ is the modality-specific feature embedding matrix whose columns correspond to embeddings of the features (image words or genes), and $\alpha_{z_{sn}} \in \mathbb{R}^D$ denotes the $D$-dimensional embedding of topic $z_{sn}$.

To fit InSTaPath to multimodal spatial transcriptomics data, we aim to maximize the log-likelihood of the observed data. Let $\mathbf{X}$ denote the collection of paired histology and gene expression observations across $S$ spatial spots:

$$\mathbf{X} := \{(\mathbf{x}_1^{(img)}, \mathbf{x}_1^{(gene)}), \ldots, (\mathbf{x}_S^{(img)}, \mathbf{x}_S^{(gene)})\}.$$

For each spot $s$, $\mathbf{x}_s^{(img)}$ and $\mathbf{x}_s^{(gene)}$ denote collections of $N_s^{(img)}$ and $N_s^{(gene)}$ words from the image and gene expression modalities, respectively. The marginal log-likelihood of the observed data is then given by:

$$\log p(\mathbf{X} \mid \alpha, \boldsymbol{\rho}) = \sum_{s=1}^{S} \log p(\mathbf{x}_s^{(img)} \mid \alpha, \rho^{(img)}) + \sum_{s=1}^{S} \log p(\mathbf{x}_s^{(gene)} \mid \alpha, \rho^{(gene)}).$$

Here, $p(\mathbf{x}_s^{(m)} \mid \alpha, \rho^{(m)})$, for $m \in \{\text{“img”}, \text{“gene”}\}$, involves an integral over the topic proportions that is intractable to compute. To address this, we reparameterize the model using the unconstrained latent variable $\delta_s$. The likelihood can then be written as:

$$p(\mathbf{x}_s^{(m)} \mid \alpha, \rho^{(m)}) = \int p(\delta_s) \prod_{n=1}^{N_s^{(m)}} p(w_{sn}^{(m)} \mid \delta_s, \alpha, \rho^{(m)})\, d\delta_s.$$

Variational inference is then used by introducing an approximate posterior distribution $Q(\delta|\, \mathbf{X}, \nu)$, where $\nu$ denotes the parameters of the encoder neural network. Assuming that $Q(\delta|\, \mathbf{X}, \nu)$ follows a simple distribution such as a Gaussian, this approximation serves as a tractable substitute for the true posterior $P(\delta|\, \mathbf{X}, \alpha, \rho)$. The inference problem can now be framed as an optimization problem by minimizing the KL divergence between these two distributions:

$$\begin{aligned}
&D_{\mathrm{KL}}[Q(\delta \mid \mathbf{X}, \nu) \,\|\, P(\delta \mid \mathbf{X}, \alpha, \boldsymbol{\rho})] \\
&= \int Q(\delta \mid \mathbf{X}, \nu) \log \frac{Q(\delta \mid \mathbf{X}, \nu)}{P(\delta \mid \mathbf{X}, \alpha, \boldsymbol{\rho})}\, d\delta \\
&= \mathbb{E}_{\delta \sim Q(\delta|\mathbf{X}, \nu)} [\log Q(\delta \mid \mathbf{X}, \nu) - \log P(\delta \mid \mathbf{X}, \alpha, \boldsymbol{\rho})] \\
&= \mathbb{E}_{\delta \sim Q(\delta|\mathbf{X}, \nu)} \left[\log Q(\delta \mid \mathbf{X}, \nu) - \log \frac{P(\mathbf{X} \mid \delta, \alpha, \boldsymbol{\rho})\, P(\delta \mid \alpha, \boldsymbol{\rho})}{P(\mathbf{X} \mid \alpha, \boldsymbol{\rho})}\right] \\
&= \mathbb{E}_{\delta \sim Q(\delta|\mathbf{X}, \nu)} [\log Q(\delta \mid \mathbf{X}, \nu) - \log P(\mathbf{X} \mid \delta, \alpha, \boldsymbol{\rho}) \\
&\qquad - \log P(\delta \mid \alpha, \boldsymbol{\rho}) + \log P(\mathbf{X} \mid \alpha, \boldsymbol{\rho})].
\end{aligned}$$

Rearranging the terms, we obtain:

$$\begin{aligned}
&\log P(\mathbf{X} \mid \alpha, \boldsymbol{\rho}) - D_{\mathrm{KL}}[Q(\delta \mid \mathbf{X}, \nu) \,\|\, P(\delta \mid \mathbf{X}, \alpha, \boldsymbol{\rho})] \\
&= \mathbb{E}_{\delta \sim Q(\delta|\mathbf{X}, \nu)} [\log P(\mathbf{X} \mid \delta, \alpha, \boldsymbol{\rho})] \\
&\quad - \mathbb{E}_{\delta \sim Q(\delta|\mathbf{X}, \nu)} [\log Q(\delta \mid \mathbf{X}, \nu) - \log P(\delta \mid \alpha, \boldsymbol{\rho})] \\
&= \mathbb{E}_{\delta \sim Q(\delta|\mathbf{X}, \nu)} [\log P(\mathbf{X} \mid \delta, \alpha, \boldsymbol{\rho})] - D_{\mathrm{KL}}[Q(\delta \mid \mathbf{X}, \nu) \,\|\, P(\delta \mid \alpha, \boldsymbol{\rho})].
\end{aligned}$$

Note that KL divergence is non-negative, and therefore, we obtain the following evidence lower bound (ELBO):

$$\begin{aligned}
\log P(\mathbf{X} \mid \alpha, \boldsymbol{\rho}) \geq & \mathbb{E}_{\delta \sim Q(\delta|\mathbf{X}, \nu)} [\log P(\mathbf{X} \mid \delta, \alpha, \boldsymbol{\rho})] \\
& - D_{\mathrm{KL}}[Q(\delta \mid \mathbf{X}, \nu) \,\|\, P(\delta \mid \alpha, \boldsymbol{\rho})] \equiv \mathrm{ELBO},
\end{aligned} \tag{1}$$

where the first term corresponds to the reconstruction likelihood, and the second term ensures that the learned distribution $Q(\delta \mid \mathbf{X}, \nu)$ remains close to the prior $\mathcal{N}(0, I_K)$ over the $K$-dimensional latent topic representation.

Model parameters are learned by maximizing the evidence lower bound (ELBO; Eq. 1), which consists of a reconstruction term and a KL divergence regularization term. The reconstruction term is defined using a multinomial distribution to model the discrete count data for each modality. In practice, we optimize the model by minimizing the negative ELBO (NELBO) using stochastic variational inference. During training, modality-specific encoders map normalized input observations to latent topic representations for each spatial spot, while the decoder reconstructs modality-specific feature distributions over genes and image words. Model parameters are updated through gradient-based optimization using minibatches of spots. The overall training procedure is summarized in Algorithm 1.

#### S3 Evaluation Metrics

Two topic modeling metrics are used to evaluate the quality of the learned topics: topic coherence (TC) and topic diversity (TD).

**Topic coherence (TC)** measures the semantic consistency of a topic by quantifying how often its top $s$ words co-occur within the same documents. It is defined as the average Normalized Pointwise Mutual Information (NPMI) over all pairs of top words within each topic, further

**Algorithm 1** Topic modeling with the InSTaPath

Initialize model and variational parameters
**for** $i = 1, 2, \ldots$ **do**
  **for** $m$ in {`image`, `gene`} **do**
    Compute $\beta_k^{(m)} = \mathrm{softmax}(\rho^{(m)\top} \alpha_k)$ for each topic $k$
  **end for**
  Choose a minibatch $B$ of spots
  **for** each spot $s$ in $B$ **do**
    **for** $m$ in {`image`, `gene`} **do**
      Get normalized bag-of-words representation $x_s^{(m)}$
      Compute $\mu_s^{(m)} = \mathrm{NN}(x_s^{(m)}; \nu_\mu^{(m)})$
      Compute $\Sigma_s^{(m)} = \mathrm{NN}(x_s^{(m)}; \nu_\Sigma^{(m)})$
    **end for**
    Compute the covariance of the joint Gaussian: $\Sigma_s^* = \left(\sum_m (\Sigma_s^{(m)})^{-1}\right)^{-1}$
    Compute the mean of the joint Gaussian: $\mu_s^* = \Sigma_s^* \left(\sum_m (\Sigma_s^{(m)})^{-1} \mu_s^{(m)}\right)$
    Sample $\theta_s \sim \mathcal{LN}(\mu_s^*, \Sigma_s^*)$
    **for** $m$ in {`image`, `gene`} **do**
      **for** each word $w_{sn}$ in the spot **do**
        Compute $p(w_{sn} \mid \theta_s) = \theta_s^\top \beta_{\cdot, w_{sn}}^{(m)}$
      **end for**
    **end for**
  **end for**
  Estimate the NELBO and its gradient (backpropagation)
  Update model parameters $\alpha, \boldsymbol{\rho}$
  Update variational parameters $(\nu)$
**end for**

averaged across all topics. Higher TC indicates that the words within a topic tend to appear together in the data, making the topic more interpretable. Topic coherence (TC) is formulated as:

$$TC = \frac{1}{K} \sum_{k=1}^{K} \frac{1}{\binom{s}{2}} \sum_{1 \le i < j \le s} \mathrm{NPMI}(w_{ki}, w_{kj}),$$

where the NPMI between the $i^{th}$ and $j^{th}$ top words from topic $k$ is

$$\mathrm{NPMI}(w_{ki}, w_{kj}) = \frac{\log\left(\dfrac{P(w_{ki}, w_{kj})}{P(w_{ki})\, P(w_{kj})}\right)}{-\log P(w_{ki}, w_{kj})}.$$

**Topic diversity (TD)** measures how different the topics are from each other by quantifying the extent to which they reuse the same top words. It is defined as the fraction of unique words among the top $s$ words across all topics. High TD indicates distinct, non-redundant topics, while low TD suggests overlapping topics with shared vocabularies. Topic diversity is formulated as:

$$TD = \frac{\left|\bigcup_{k=1}^{K} \mathcal{W}_k\right|}{K\, s}, \qquad \mathcal{W}_k = \{w_{k1}, w_{k2}, \ldots, w_{ks}\}.$$

Three clustering metrics are used to evaluate the quality of spatial domain identification: Adjusted Rand Index (ARI), Normalized Mutual Information (NMI), and Average Silhouette Width (ASW). These metrics quantify the agreement between predicted cluster assignments and ground-truth annotations, as well as the compactness and separation of the learned embeddings. All metrics were computed using the `scikit-learn` implementation (`adjusted_rand_score`, `normalized_mutual_info_score`, and `silhouette_score`).

**Adjusted Rand Index (ARI).** ARI measures the similarity between two clusterings by considering all pairs of samples and counting pairs that are assigned consistently in both clusterings. Random labeling has an ARI close to 0. An ARI of 1 indicates perfect agreement between the predicted clusters and the ground truth.

**Normalized Mutual Information (NMI).** NMI quantifies the shared information between predicted clusters and ground-truth labels. NMI ranges from 0 (no mutual information) to 1 (perfect correlation).

**Average Silhouette Width (ASW).** ASW evaluates the compactness and separation of clusters based on pairwise distances in the embedding space. The original ASW score lies in the range $[-1, 1]$, where higher values indicate better cluster separation. For ease of comparison across metrics, we linearly rescale the ASW score to the range $[0, 1]$.

#### S4 Related work

**STAMP** is a spatially aware topic modeling framework for spatial transcriptomics that models each spot as a mixture of latent topics representing gene expression programs. The method combines topic modeling with a simplified graph convolutional network (SGCN) to incorporate spatial relationships between neighboring spots. STAMP outputs spatial topics and associated gene modules, enabling interpretable identification of spatial domains and gene signatures without explicit clustering.

**OmiCLIP** is a multimodal framework that aligns histology images with transcriptomic profiles using CLIP-style contrastive learning. The model takes as input spot-level histology image patches together with the corresponding gene expression profiles. The image encoder is based on a vision transformer (ViT) operating on $224 \times 224$ pixel image patches, while the gene modality is encoded using a causal masking transformer. To represent transcriptomic information, gene expression profiles are converted into "sentences" by concatenating the top 50 expressed genes for each spot.

#### S5 Evaluation and robustness of ViT-VQ image-word representations

To evaluate the representation quality of the ViT-VQ image-word features, we compared them with the original UNI image embeddings using image patches from the CRC-100k/CRC-7K datasets. Figure S1 shows the t-SNE projections of both representations. Both methods separate image patches from different tissue classes well. However, UNI features tend to produce more scattered clusters for patches belonging to the same tissue class, likely reflecting batch effects across different patients. To examine this, we colored the t-SNE plots by dataset source (CRC-100k vs CRC-7K). Although patient IDs are unavailable, these two datasets are known to originate from different WSIs of non-overlapping patients. The results show that ViT-VQ features achieve better mixing of points from the two datasets for most tissue types, suggesting reduced batch effects. This observation is further supported by the higher kBET score reported in Table S1. In addition, ViT-VQ representations achieve improved clustering performance in terms of ARI, NMI, and ASW.

Because ViT-VQ converts continuous embeddings into count-based image words through vector quantization, some fine-grained information is inevitably compressed. Therefore, we expect the raw UNI features to achieve higher classification accuracy in linear probing. To test this, we evaluated both representations using multiclass AUROC under a one-vs-one (OvO) scheme, where binary classifiers are trained for each pair of classes and their AUROC scores are averaged (Table S1). Interestingly, ViT-VQ achieves slightly higher AUROC than the raw UNI features, which may be attributed to its more compact and unified cluster representation.

Finally, we tested whether the grouping of feature dimensions affects the ViT-VQ representation. Specifically, we randomly permuted the 1536-dimensional UNI image features across dimensions before the vector quantization step, using 100 random seeds to generate different feature groupings. The ViT-VQ pipeline and linear probing evaluation were then applied to each permuted representation. Experiments were conducted on a class-balanced subset of the CRC-100k/CRC-7K datasets, with 1,000 patches per tissue class used for training and 100 patches per class for testing. As shown in Table S2 and Figure S2, random regrouping of image feature dimensions has minimal impact on downstream classification performance. These results indicate that the ViT-VQ representation is robust to the grouping of feature dimensions, supporting the use of the original UNI output and partitioning based on the attention head dimension.

#### Supplementary Tables

**Table S1** Linear probe, clustering, and batch effect evaluation results on the CRC-100K/CRC-7K dataset.

| | Linear Probe | | | | | Clustering | | | Batch effect |
|---|---|---|---|---|---|---|---|---|---|
| Method | ACC | BACC | KAPPA | F1 | AUROC | ARI | NMI | ASW | kBET |
| UNI | 0.969 | 0.957 | 0.988 | 0.969 | 0.989 | 0.16 | 0.52 | 0.54 | 0.43 |
| ViT-VQ | 0.910 | 0.901 | 0.981 | 0.913 | 0.992 | 0.48 | 0.72 | 0.67 | 0.58 |

ACC: Accuracy; BACC: Balanced Accuracy; KAPPA: Cohen's $\kappa$; F1: weighted-average F1 score; AUROC: multi-class AUROC computed with the one-vs-one (OvO) scheme. ARI: Adjusted Rand Index; NMI: Normalized Mutual Information; ASW: Average Silhouette Width; kBET: k-nearest neighbor Batch Effect Test. ARI and NMI were computed from cluster assignments obtained using the `mclust` R package, with the number of clusters set to match the ground truth. All metrics are reported such that higher values indicate better performance. Cohen's $\kappa$ takes values in $[-1, 1]$, while all other metrics are bounded in $[0, 1]$.

**Table S2** Linear probe performance comparison.

| Method | ACC | BACC | KAPPA | F1 | AUROC |
|---|---|---|---|---|---|
| UNI | 0.968 | 0.968 | 0.991 | 0.968 | 0.992 |
| ViT-VQ | 0.948 | 0.948 | 0.985 | 0.947 | 0.995 |
| ViT-VQ (100 seeds)[1] | 0.949<br>$\pm 0.016$ | 0.949<br>$\pm 0.016$ | 0.975<br>$\pm 0.008$ | 0.945<br>$\pm 0.017$ | 0.996<br>$\pm 0.001$ |

All metrics are reported as higher-is-better.
[1] Results are reported as mean $\pm$ standard deviation over 100 random seeds.

### Supplementary Figures

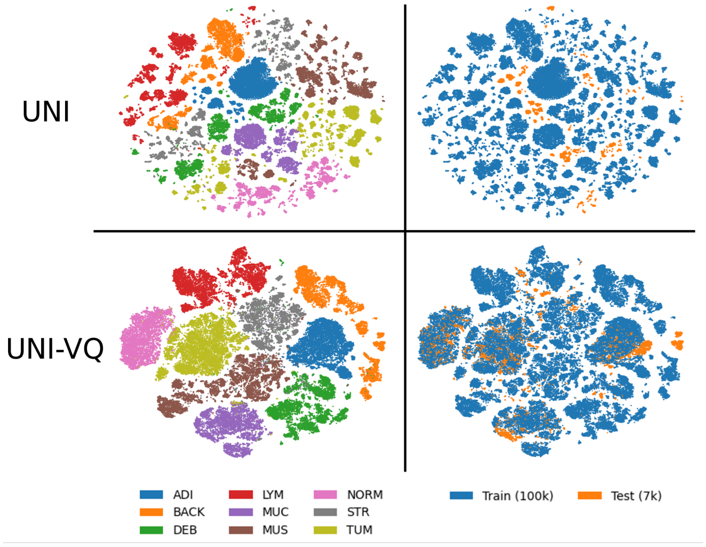


**Figure S1** t-SNE projections of image patches from the CRC-100k/CRC-7K datasets using UNI image features (top) and ViT-VQ image word count representations (bottom). Left panels are colored by tissue class, showing that both representations separate tissue types well, while right panels are colored by dataset (CRC-100k vs CRC-7K), illustrating improved mixing across datasets with ViT-VQ compared to the raw UNI features.

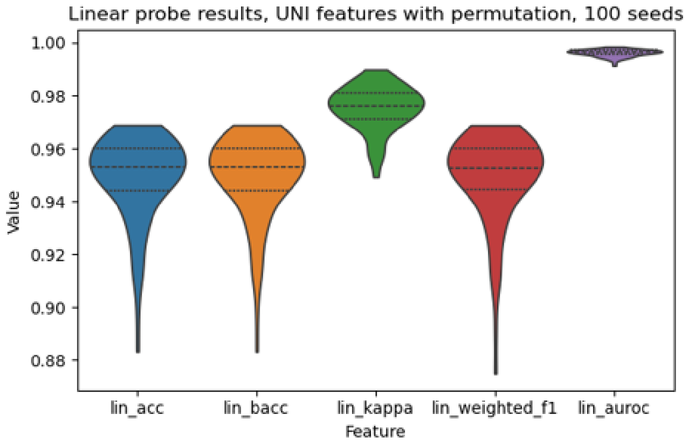


**Figure S2** Violin plots show the distribution of linear probing performance across 100 random seeds after randomly permuting the 1536-dimensional UNI image features before the VQ step in the ViT-VQ pipeline. Evaluation on a class-balanced subset of the CRC-100k/CRC-7K datasets shows that random regrouping of image feature dimensions has minimal impact on downstream classification performance.

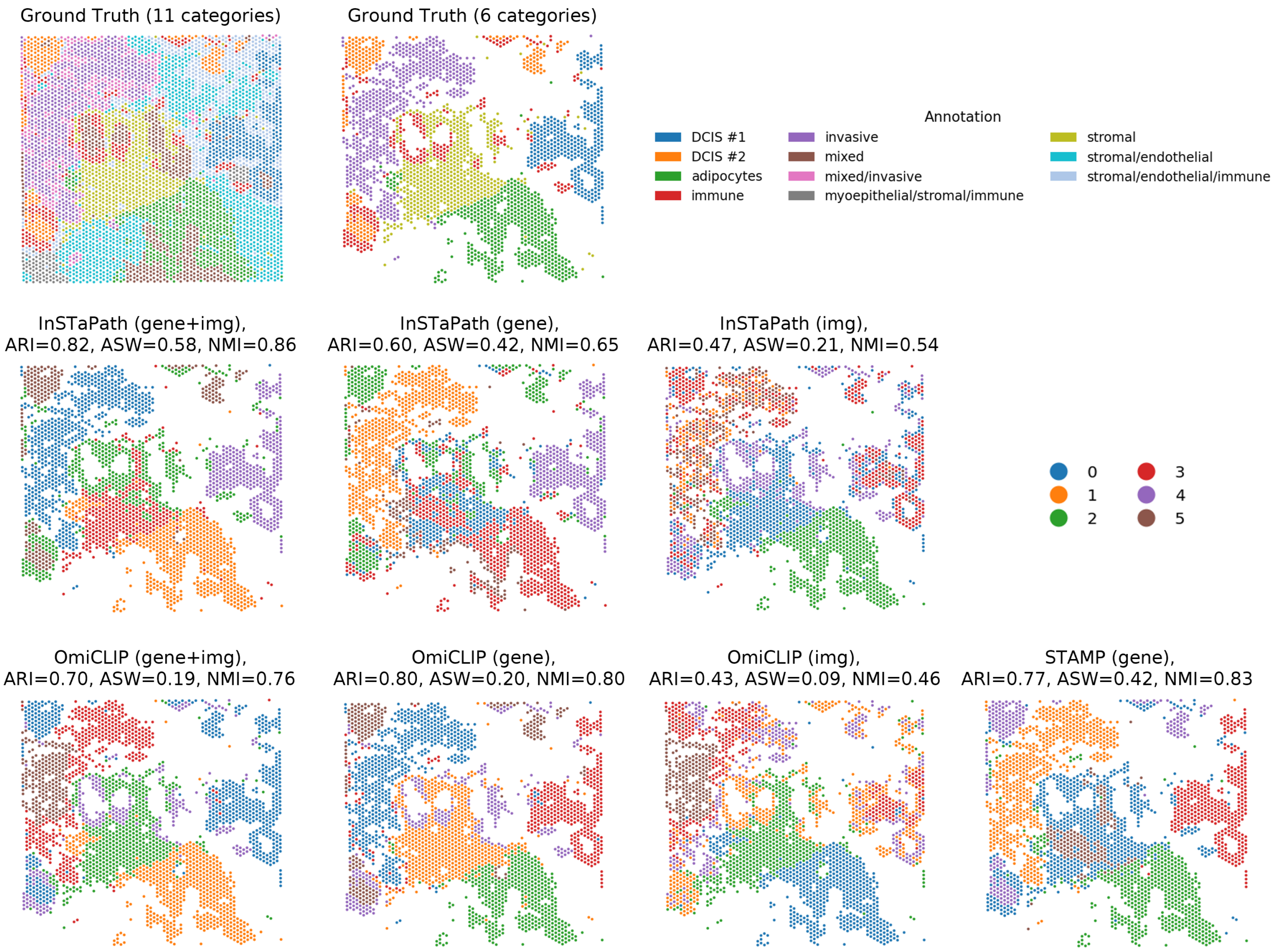


**Figure S3** Benchmark comparison of spot-level embeddings across methods. Clustering results are shown as the raw output from the `mclust` R package without additional smoothing or refinement.

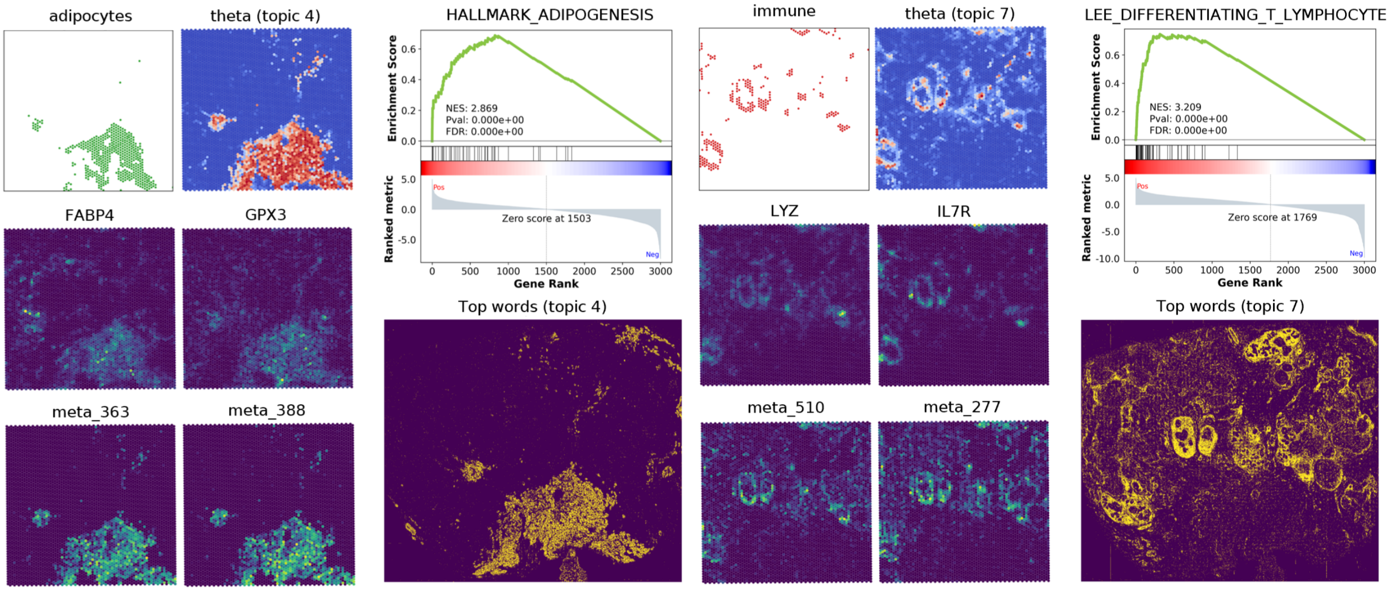


**Figure S4** Additional examples related to Fig. 3d, illustrating the spatial alignment between representative genes and image words, along with GSEA results and whole-slide distributions of the corresponding top image words.

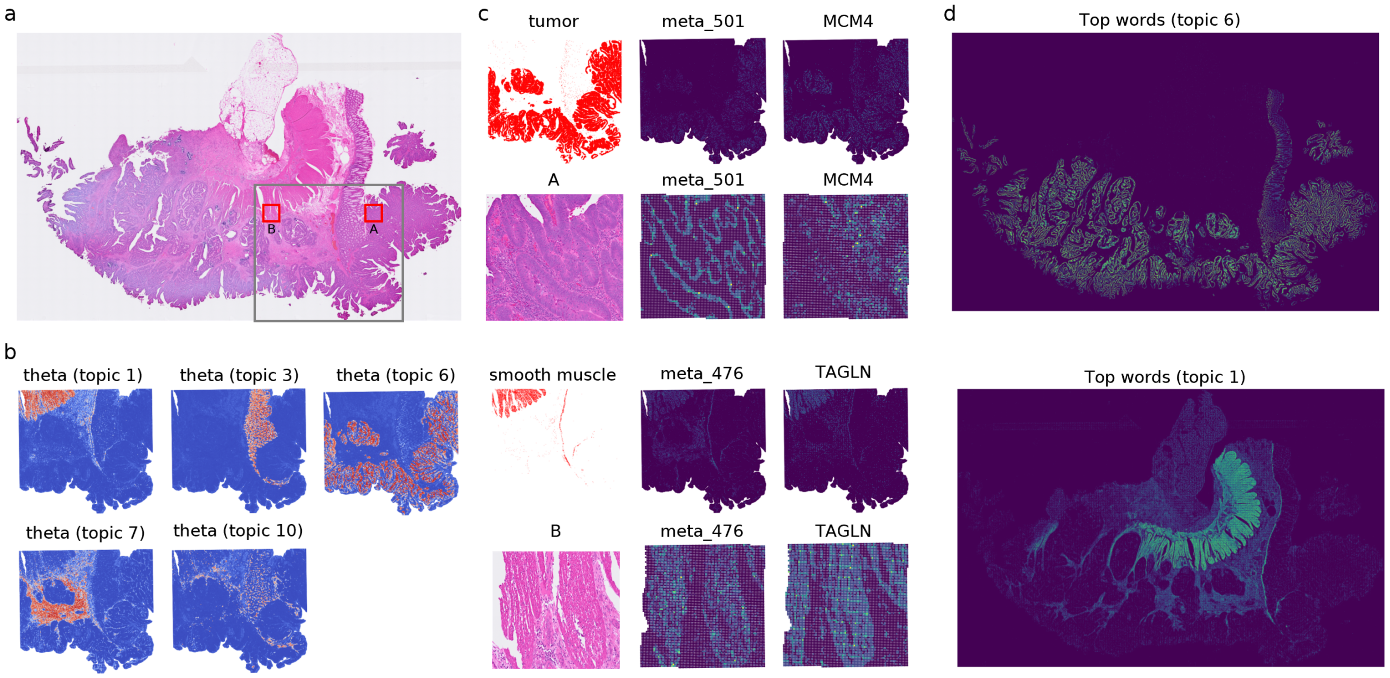


**Figure S5** **(a)** Whole-slide histology image. The gray box indicates the region used for spatial transcriptomics profiling, and the red box marks the region of interest (ROI). **(b)** Spatial distribution of spot–topic proportions across the tissue section. Each spot is colored according to the inferred proportion of a given topic. **(c)** Examples of tumor-associated Topic 6 and smooth muscle–associated Topic 1 within the zoomed-in ROI, showing the spatial distribution of representative image words and gene expression. **(d)** Distribution of the top image words across the full histology image.
